# Glycosylphosphatidylinositol biosynthesis restricts coronavirus infection via the regulation of LY6E

**DOI:** 10.1101/2025.02.08.637211

**Authors:** Yanlong Ma, Fei Feng, Hui Feng, Xue Ma, Ziqiao Wang, Yutong Han, Yunkai Zhu, Yuyan Wang, Zhichao Gao, Yuyuan Zhang, Jincun Zhao, Rong Zhang

**Affiliations:** Key Laboratory of Medical Molecular Virology (MOE/NHC/CAMS), Shanghai Institute of Infectious Disease and Biosecurity, Shanghai Frontiers Science Center of Pathogenic Microorganisms and Infection, School of Basic Medical Sciences, Shanghai Medical College, Fudan University, Shanghai 200032, China; State Key Laboratory of Respiratory Disease, National Clinical Research Center for Respiratory Disease, Guangzhou Institute of Respiratory Health, the First Affiliated Hospital of Guangzhou Medical University, Guangzhou, Guangzhou 510182, China

**Keywords:** Coronavirus, CRISPR screening, GPI, GPI-anchored proteins, LY6E

## Abstract

Coronaviruses, including SARS-CoV-2, rely on host factors for their replication and pathogenesis, while hosts deploy defense mechanisms to counteract viral infections. Although numerous host proviral factors have been identified, the landscape of host restriction factors and their underlying mechanisms remain less explored. Here, we conducted genome-wide CRISPR knockout screens using three distinct coronaviruses—SARS-CoV-2, HCoV-OC43 (a common cold human virus from the genus *Betacoronavirus*) and porcine epidemic diarrhea virus (*Alphacoronavirus*) to identify conserved host restriction factors. We identified glycosylphosphatidylinositol (GPI) biosynthesis as the pan-coronavirus host factor that restrict viral entry by disrupting spike protein-mediated membrane fusion at both endosomal and plasma membranes. GPI biosynthesis generates GPI moieties that covalently anchor proteins (GPI-anchored proteins [GPI-APs]) to the cell membrane, playing essential roles in various cellular processes. Through focused CRISPR knockout screens targeting 193 GPI-APs, we identified LY6E as the key downstream effector mediating the antiviral activity of the GPI biosynthesis pathway. These findings reveal a novel role for GPI biosynthesis as a conserved host defense mechanism against coronaviruses and highlight LY6E as a critical antiviral effector. This study provides new insights into virus-host interactions and the development of host-directed antiviral therapies.

## INTRODUCTION

*Coronaviridae* is a family of enveloped, positive-sense, single-stranded RNA viruses and consists of four genera: *Alphacoronavirus*, *Betacoronavirus*, *Gammacoronavirus*, and *Deltacoronavirus*. Coronaviruses exhibit broad host tropism, infecting various species, including humans, pigs, birds, cattle, dogs, and cats. Although many human coronaviruses, (e.g., HCoV-OC43, HCoV-229E, HCoV-NL63, and HCoV-HKU1) typically cause mild, cold-like symptoms^1, 2^, others, such as SARS-CoV, MERS-CoV, and SARS-CoV-2, can result in severe respiratory syndromes with high morbidity and mortality rates. In particular, SARS-CoV-2, the causative agent of the COVID-19 pandemic, has led to over 600 million confirmed cases and nearly 7 million deaths worldwide^3^. Numerous SARS-CoV-2 variants, including alpha, beta, delta, and omicron, have emerged over the course of the pandemic and continue to evolve, posing significant threats to public health^4^.

Coronaviruses infecting animals in close contact with humans, such as livestock, can significantly impact on the food industry. Transmissible gastroenteritis virus, porcine epidemic diarrhea virus (PEDV), swine acute diarrhea syndrome coronavirus, and porcine deltacoronavirus (PDCoV) are coronaviruses that can cause severe diarrhea, vomiting, and dehydration in pigs, particularly piglets^5, 6^. In poultry, multiple serotypes and strains of the infectious bronchitis virus (IBV) varying tissue tropism and pathogenicity have emerged, causing extensive damage to the poultry industry^7^. Additionally, some of these viruses exhibit zoonotic potential. For example, PDCoV has occasionally infected humans^8^. Human coronaviruses like SARS-CoV, MERS-CoV, SARS-CoV-2, HCoV-NL63, and HCoV-229E, are believed to have originated from bats^9, 10^. A human coronavirus identified in 2021, CCoV-HuPn-2018, is thought to have originated from canine coronaviruses^11^. Coronaviruses mutate and recombine to adapt to new hosts, expanding their host range and tissue tropism. Thus, a deeper understanding of their biology, pathogenesis, and host interactions is crucial for global health.

Coronaviruses rely on host factors for entry, replication, assembly, and release, while hosts activate immune defenses that viruses counteract through evasion strategies^2^. Understanding these interactions is vital for the development of host-directed antiviral therapies. Unbiased genetic screens are powerful tools for studying virus-host interactions^12, 13^. Following the COVID-19 pandemic, many CRISPR-based loss- and gain-of-function screens have identified key proviral host factors for SARS-CoV-2 infection, including ACE2, TMPRSS2, and CTSL, as well as other factors like TMEM41B, TMEM106B, HMGB1, SCAP, RAB7A, class III PI3K subunits, and SWI/SNF complex^14, 15, 16, 17, 18, 19, 20^. However, most of these screens have focused on proviral factors, with few exploring host restriction factors. Studies using interferon-stimulated gene (ISG)–based cDNA libraries or CRISPR knockout/activation screens have identified ISGs such as LY6E, BST2, DAXX, OAS1, and DDX41 as restriction factors for coronaviruses^21, 22, 23^. Beyond ISGs, whole-genome CRISPR screens have identified mucins and other broad-spectrum antiviral factors like CH25H, ZAP, LARP1, PLSCR1, and DAZAP2^24, 25, 26, 27, 28^.

Given the conserved replication strategy of coronaviruses^2^, in the present study, we conducted whole-genome CRISPR knockout screens using three distinct coronaviruses: SARS-CoV-2, HCoV-OC43 (*Betacoronavirus*), and PEDV (*Alphacoronavirus*). We identified glycosylphosphatidylinositol (GPI) biosynthesis genes as pan-coronavirus restriction factors that inhibit spike-mediated endolysosomal and plasma membrane fusion. GPI biosynthesis, essential for embryogenesis and immune responses, generates GPI moieties that covalently anchor various proteins (known as GPI-anchored proteins [GPI-AP]) to the cell membrane^29, 30, 31^. Through focused CRISPR screens, we specifically identified one of these GPI-APs, LY6E, as a key downstream anti-coronavirus effector of the GPI biosynthesis pathway. These findings reveal a novel role for GPI biosynthesis as a conserved host defense mechanism against coronaviruses.

## RESULTS

### Genome-wide CRISPR/Cas9 knockout screens identify host restriction factors for coronavirus infection

To identify pan-coronavirus restriction factors, we performed genome-wide, cell sorting–based screens in A549 cells expressing the receptor ACE2 (A549-ACE2) using three coronaviruses: a. transcription- and replication-competent SARS-CoV-2 virus-like particles (SARS-CoV-2 trVLP-GFP), in which the nucleocapsid (N) gene is replaced by the GFP reporter gene and the particles are trans-packaged in N-expressing cells^32^; b. HCoV-OC43-mGreen, in which the mGreenLantern reporter gene was inserted into the HCoV-OC43 genome in place of the ns2a gene; and c. PEDV-GFP, in which the GFP is inserted into the PEDV genome in place of ORF3. SARS-CoV-2 (original strain) and HCoV-OC43, both *Betacoronavirus*, primarily infect the lower and upper respiratory tracts in humans, respectively, while PEDV, an *Alphacoronavirus*, infects the intestines of pigs but can also infect human cell lines. The screening in A549-ACE2 cells with SARS-CoV-2 trVLP-GFP was similar to a previous study^28^ but conducted at a lower MOI to reduce the signal-to-noise ratio. To rule out cell type–specific effects, HeLa cells containing the genome-wide CRISPR knockout library were also infected with PEDV-GFP. Knockout of genes with antiviral activity would presumably enhance virus infection, so we sorted the virus-infected, reporter-positive cells to enrich susceptible cells for genomic DNA extraction, sgRNA sequencing, and data analysis (**Fig. 1A** and **Supplementary Table 1)**.

**Fig. 1.**
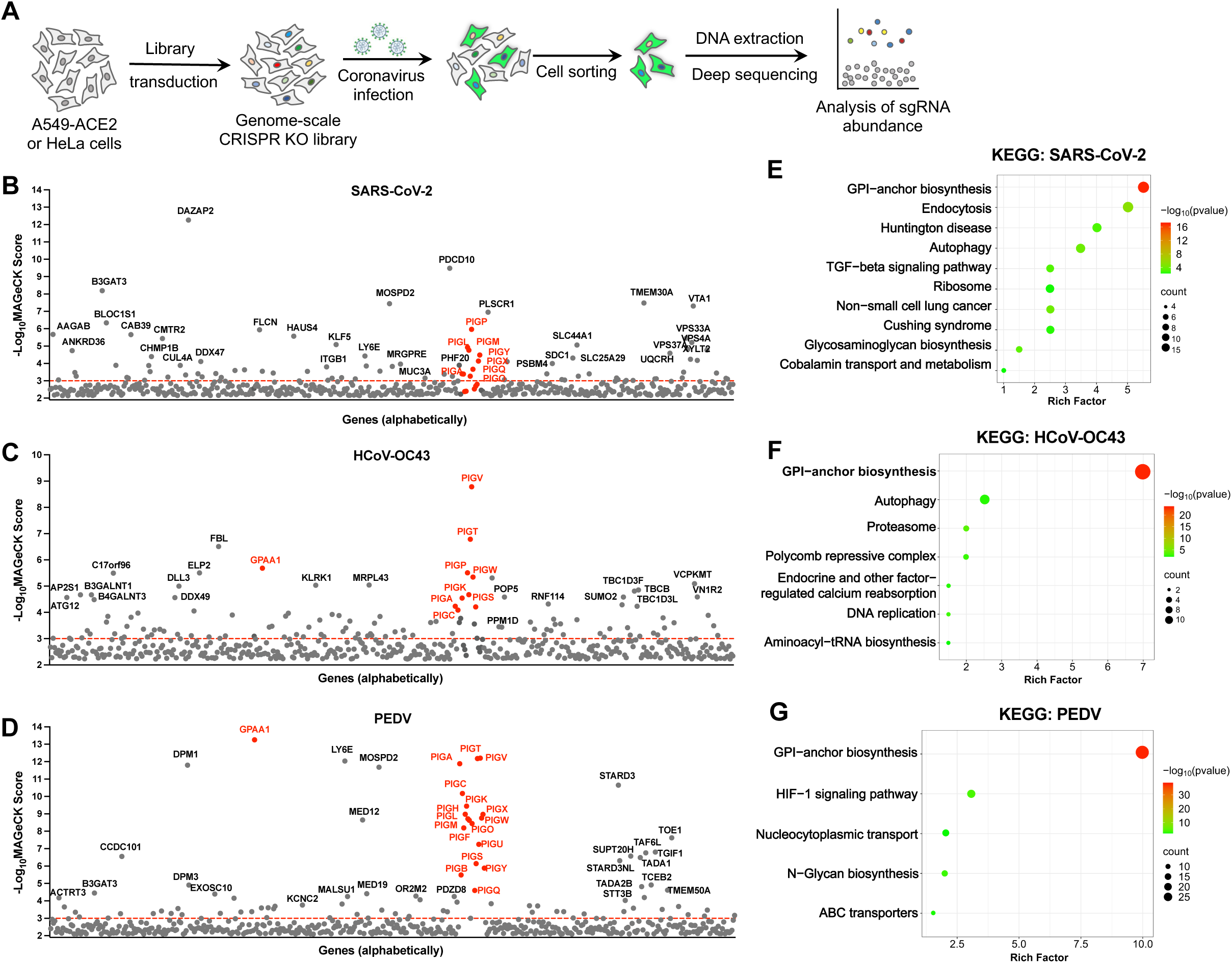
Genome-wide CRISPR/Cas9 knockout screens identify host restriction factors for coronavirus infection. **A.** Workflow of the genome-wide CRISPR screening. A549-ACE2 or HeLa cells expressing the Cas9 were transduced with a CRISPR knockout sgRNA library, followed by infection with coronaviruses expressing fluorescent protein reporter. Infected reporter-positive cells were sorted for genomic extraction and sgRNA sequence analysis. **B-D.** Genes identified from the CRISPR screens in A549-ACE cells using SARS-CoV-2 trVLP-GFP **(B)**, HCoV-OC43-mGreen **(C)**, or PEDV-GFP **(D)** at an MOI of 0.1 for 24 h. The genes were analyzed by MAGeCK software and sorted based on the -log_10_(MAGeCK score). **E-G.** KEGG pathway analysis of 100 top-ranked genes from the screens in **B-D**.

The genes identified by the screens were ranked by MAGeCK score. Among the candidate genes, *DAZAP2* and *PLSCR1* were identified in the SARS-CoV-2 screen in A549-ACE2 cells, while *LY6E* and *MOSPD2* were found in both the SARS-CoV-2 and PEDV screens in A549-ACE2 or HeLa cells **(Fig. 1B** and **D; Supplementary Fig. 1A)**. *DAZAP2*^27, 28^, *PLSCR1*^26^, *LY6E*^33, 34^ and *MOSPD2*^16^ have previously been reported as important host restriction factors against coronaviruses, supporting the validity of our screens. Interestingly, GPI biosynthesis genes, including *PIGA, PIGP, PIGV, PIGO, PIGX, PIGM,* and *GPAA1*, were significantly enriched in the screens of all three coronaviruses **(Fig. 1B-D; Supplementary Fig. 1A)**. KEGG analysis of the top 100 genes showed that most are involved in GPI-anchor biosynthesis pathway **(Fig. 1E-G)**.

### Identification of genes involved in GPI biosynthesis as pan-coronavirus host restriction factors

To cross-validate and identify conserved hits, we selected the 50 top-ranked genes from the PEDV screen in HeLa cells **(Supplementary Fig. 1A)**. Knockout HeLa cell lines were constructed using two sgRNAs for each of the 50 genes, and PEDV infection efficiency was subsequently determined by measuring the percentage of N-positive cells. Among the genes with the greatest positive effect on infection efficiency after knockout were multiple GPI biosynthesis genes: *GPAA1, PIGA, PIGV*, and *DPM1* knockouts increased infection over 8-fold compared to control, while *PIGO* knockout increased infection approximately 4-fold **(Supplementary Fig. 1B)**. Other genes that increased infection efficiency by ≥4-fold upon knockout included *MOSPD2, P4HB*, and *LY6E* (each leading to an approximately 4-fold increase) and *IRX2* (an approximately 8-fold increase). These findings suggest that GPI biosynthesis genes play key roles in restricting coronavirus infection.

In eukaryotes, GPI moieties produced by GPI biosynthesis are a form of post-translational modification that covalently link to specific proteins, anchoring them to the cell surface. Approximately 30 genes are involved in the GPI biosynthesis pathway^30,35^ **(Fig. 2A)**. In our CRISPR knockout screens in A549-ACE2 or HeLa cells using SARS-CoV-2, HCoV-OC43, and PEDV, around 20 GPI biosynthesis genes were identified. For further study, we selected *PIGA, PIGV*, and *GPAA1* from the early, intermediate, and protein-anchoring steps, respectively, of the GPI biosynthesis pathway, based on their considerable antiviral activity during PEDV infection **(Fig. 2A, Supplementary Fig. 1B)**. We first validated the antiviral activity of these three genes in the context of authentic SARS-CoV-2 infection. Knockout of *PIGA, PIGV,* and *GPAA1* in A549-ACE2 cells significantly increased SARS-CoV-2 infection, with a 3-fold increase for the original and Delta variants and a 5-7–fold increase for the omicron variants BA.1 and BA.2 **(Fig. 2B-F)**.

**Fig. 2.**
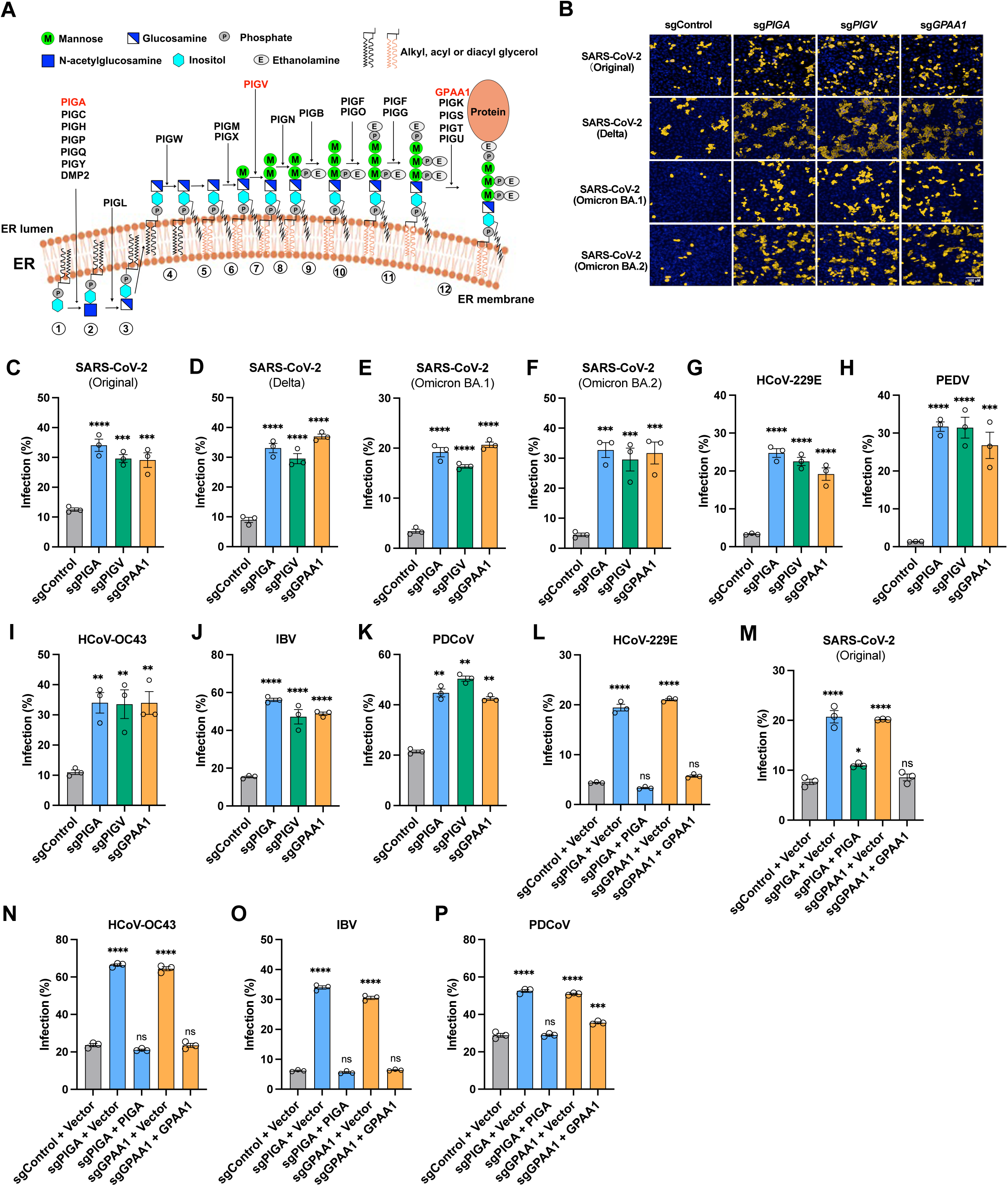
Identification of genes involved in GPI biosynthesis as pan-coronavirus host restriction factors. **A.** Schematic of the genes involved in the GPI biosynthesis pathway. **B.** Representative immunofluorescence images showing the infectivity of SARS-CoV-2 original strain and its variants in gene-knockout A549-ACE2 cells. Scale bar, 100 μm. **C-F.** High content imaging and quantification analysis of the infection by SARS-CoV-2 original **(C)**, Delta **(D)**, Omicron BA.1 **(E)**, or Omicron BA.2 **(F)** in gene-knockout A549-ACE2 cells (MOI 0.5, 24 h). **G-K.** Flow cytometry analysis of the infection by other coronaviruses. Gene-knockout HeLa cells were infected with the alphacoronavirus HCoV-229E (MOI 0.75, 32 h) **(G)** or PEDV (MOI 1, 20 h) **(H)**; the betacoronavirus HCoV-OC43 (MOI 0.5, 20 h) **(I)**; the gammacoronavirus IBV (MOI 0.75, 20 h) **(J)**; or the deltacoronavirus PDCoV (MOI 0.25, 17 h) (**K**). **L-P.** Virus infectivity in knockout cells trans-complemented with the respective genes. Flow cytometry analysis of HeLa cells infected with HCoV-229E (MOI 0.75, 32 h) **(L)**; A549-ACE2 cells infected with SARS-CoV-2 original strain (MOI 0.5, 24 h) **(M)**; HeLa cells infected with HCoV-OC43 (MOI 0.5, 20 h) **(N)**; IBV (MOI 0.75, 20 h) **(O)**; or PDCoV (MOI 0.25, 17 h) **(P)**. Data shown are from three independent experiments. One-way ANOVA with Dunnett’s test; mean ± s.d.; *P < 0.05; **P < 0.01; ***, P < 0.001; ****P < 0.0001; ns, not significant.

To investigate whether GPI biosynthesis genes represent a common host restriction pathway for the family *Coronaviridae*, we infected *PIGA-, PIGV-,* or *GPAA1*-knockout HeLa cells with viruses from the four *Coronaviridae* genera: the alphacoronaviruses HCoV-229E and PEDV; the betacoronavirus HCoV-OC43; the gammacoronavirus IBV; and the deltacoronavirus PDCoV. Knockout of each gene was verified by Inference of CRISPR Edits (ICE) analysis^36^ **(Supplementary Fig. 2)**. Notably, knockout of *PIGA, PIGV*, or *GPAA1* significantly increased the infection of other coronaviruses relative to controls, with the strongest effects observed for HCoV-229E (7-fold) and PEDV (20-fold) **(Fig. 2G** and **H)**. The infection efficiency of HCoV-OC43, IBV, and PDCoV increased 2-4 fold **(Fig. 2I-K)**.

Addtionally, the antiviral function in *PIGA*- and *GPAA1*-knockout cells was rescued by genetic complementation. We re-introduced *PIGA* or *GPAA1* cDNA into the respective knockout HeLa cells **(Supplementary Fig. 3A)** and found that trans-complementation not only significantly reduced infection with alphacoronaviruses HCoV-229E and PEDV (**Fig. 2L, Supplementary Fig. 3B**) but also with the SARS-CoV-2 original, Delta, Omicron BA.1 and BA.2 variants (**Fig. 2M, Supplementary Fig. 3C-F**). Infections with HCoV-OC43, IBV, and PDCoV were also decreased **(Fig. 2N-P)**. However, overexpression of *PIGA* or *PIGV* in unmodified HeLa cells did not alter infection by multiple coronaviruses relative to controls (**Supplementary Fig. 4**). These findings suggest that basal expression of GPI biosynthesis genes is sufficient to exhibit broad antiviral activity against different coronaviruses.

### GPI biosynthesis genes inhibit coronavirus entry

Since the downstream proteins modified by GPI biosynthesis are cell surface-anchored proteins, we hypothesized that knockout of *PIGA, PIGV*, or *GPAA1* might affect viral entry. We infected our knockout cells with VSV-based pseudovirus bearing the spike protein of the SARS-CoV-2 original strain or, as a control, the glycoprotein of VSV (VSV-G). Knockout of *PIGA, PIGV*, or *GPAA1* in A549-ACE cells significantly enhanced SARS-CoV-2 pseudovirus infection by approximately 3-fold whereas no difference was observed for VSV-G pseudovirus **(Fig. 3A** and **B)**. Moreover, knockout of these genes also markedly increased the infection by VSV-based pseudoviruses expressing the spike protein of SARS-CoV-1, HCoV-229E, HCoV-OC43, or PEDV **(Fig. 3C-F)**. These results suggest that these three genes involved in GPI biosynthesis might specifically inhibit coronavirus entry.

**Fig. 3.**
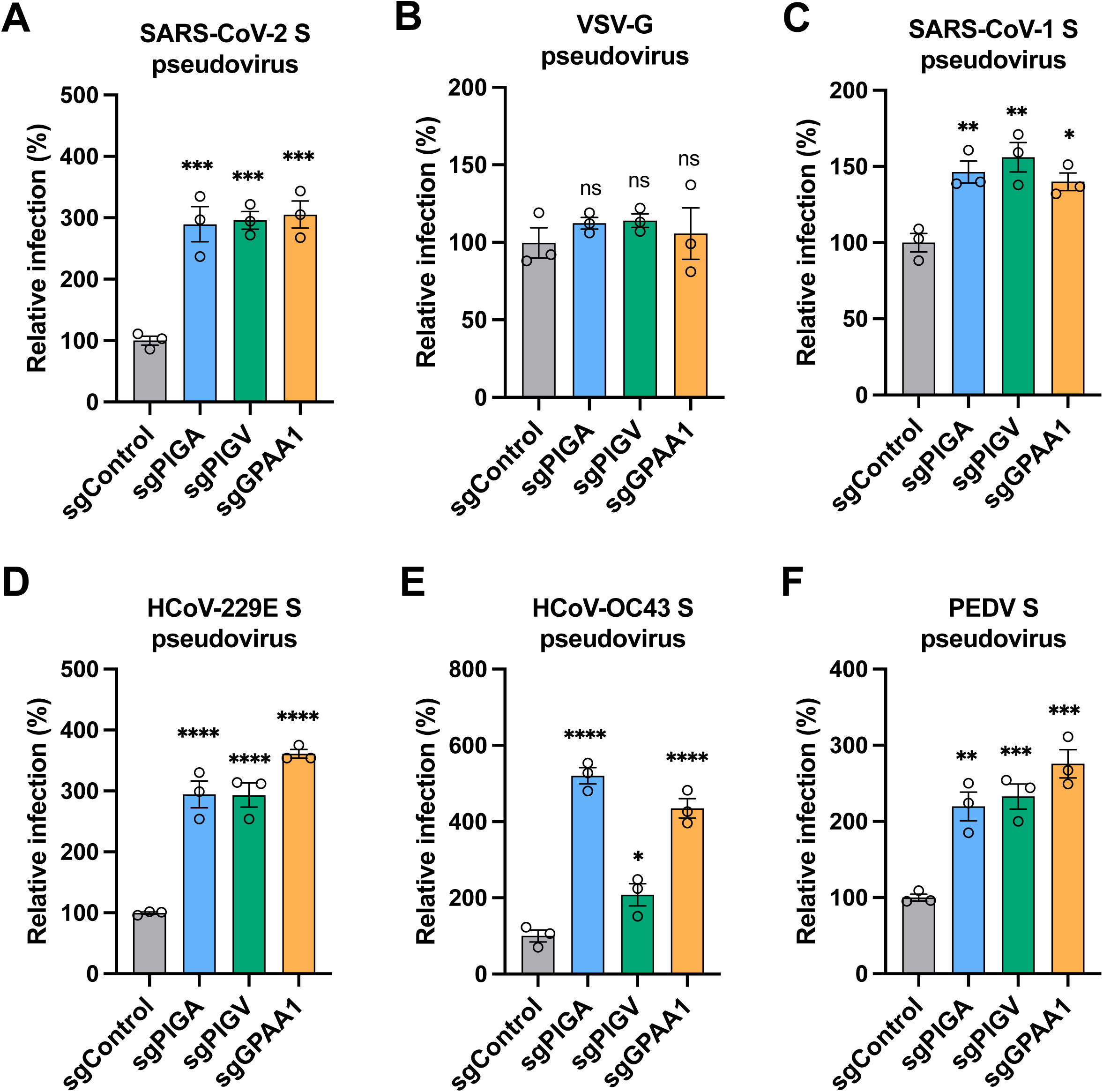
GPI biosynthesis genes inhibit coronavirus entry. **A-F.** Control, *PIGA*-, *PIGV-*, and *GPAA1*-knockout A549-ACE2 cells were infected with VSV-based pseudoviruses bearing the glycoprotein of SARS-CoV-2 original strain **(A)**, VSV **(B)**, SARS-CoV-1 **(C)**, HCoV-229E **(D)**, HCoV-OC43 **(E)**, or PEDV **(F)**, and the luciferase activity was measured and normalized to the control. Data shown are from at least three independent experiments. one-way ANOVA with Dunnett’s test; mean ± s.d.; *P < 0.05; **P < 0.01; ***, P < 0.001; ****P < 0.0001.

### GPI biosynthesis genes inhibit coronavirus entry by disrupting spike-mediated endolysosomal fusion

A549-ACE2 cells express minimal or no serine proteases such as TMPRSS2 (required for plasma membrane entry) and high level of cysteine proteases such as CTSL (required for endosomal entry). To determine whether the GPI biosynthesis affects the endosomal pathway of SARS-CoV-2 entry in A549 cells, we selected the representative *GPAA1* gene for further study. Control and *GPAA1-*knockout cells were infected with murine leukemia retrovirus (MLV)-based pseudovirus bearing the spike protein of SARS-CoV-2 in the presence of E-64d (aloxistatin), an inhibitor of the cysteine proteases, and/or camostat mesylate, an inhibitor of serine proteinases. As expected, the enhancement of virus infection in *GPAA1*-knockout cells was significantly abolished with the E-64d inhibitor, whereas camostat had no effect **(Fig. 3A)**. Similar results were obtained when cells were infected with single-round SARS-CoV-2 trVLP-NLuc particles^28^, in which the N gene is replaced by NanoLuc luciferase **(Fig. 3B)**. These results demonstrate that GPI biosynthesis pathway impacts the endosomal entry.

We next investigated the stages of endosomal entry at which the GPI biosynthesis pathway acts, using SARS-CoV-2 as the representative virus. The endosomal entry of coronavirus involves virion binding, internalization (uptake), trafficking (transport to the late endosome or lysosome), low pH-dependent activation of cysteine proteases to cleave the spike protein, spike-mediated virion fusion with endolysosome membranes, and finally viral genome release into the cytoplasm^2^. After incubating virions with control or *GPAA1*-knockout cells on ice to allow binding, or shifting to 37°C for internalization, we found that knockout of *GPAA1* did not affect virion binding or internalization **(Fig. 3C** and **D)**. Virion trafficking was examined by analyzing the co-localization of spike- and N-positive foci (virions) with the lysosome marker LAMP1. The E-64d inhibitor was used to block virion fusion with endolysosome membranes. knockout of *GPAA1* did not impact the number of virions trafficking to endolysosomes post-internalization **(Fig. 3E**, and **Supplementary Fig. 5)**. Moreover, we did not detect significant differences in endosome/lysosome acidification in *GPAA1*-knockout compared to controls, as measured using LysoSensor Green dye **(Fig. 3F)**. Consistently, cleavage of the spike protein mediated by low pH-activated cysteine proteases was similar between knockout and control cells **(Fig. 3G)**. The E-64d inhibitor was used as a control to prevent the cleavage **(Fig. 3G)**.

To investigate whether GPI biosynthesis genes regulate the fusion of viral particles with endolysosomal membranes, we used a modified split-NanoLuc system to assess virus-cell fusion^26^. Given that cyclophilin A interacts with retroviral Gag protein, the CypA-HiBiT fusion protein was packaged into MLV–based SARS-CoV-2 pseudovirus particles. Control or *GPAA1*-knockout A549-ACE2 cells stably expressing the LgBiT were infected with the pseudovirus. The NanoLuc luciferase signal is only produced when its two subunits (HiBiT and LgBiT) are reconstituted in the same cell. We found that *GPAA1* knockout significantly increased the luciferase signal, indicating enhanced virion fusion with endolysosomal membranes **(Fig. 4H)**.

**Fig. 4.**
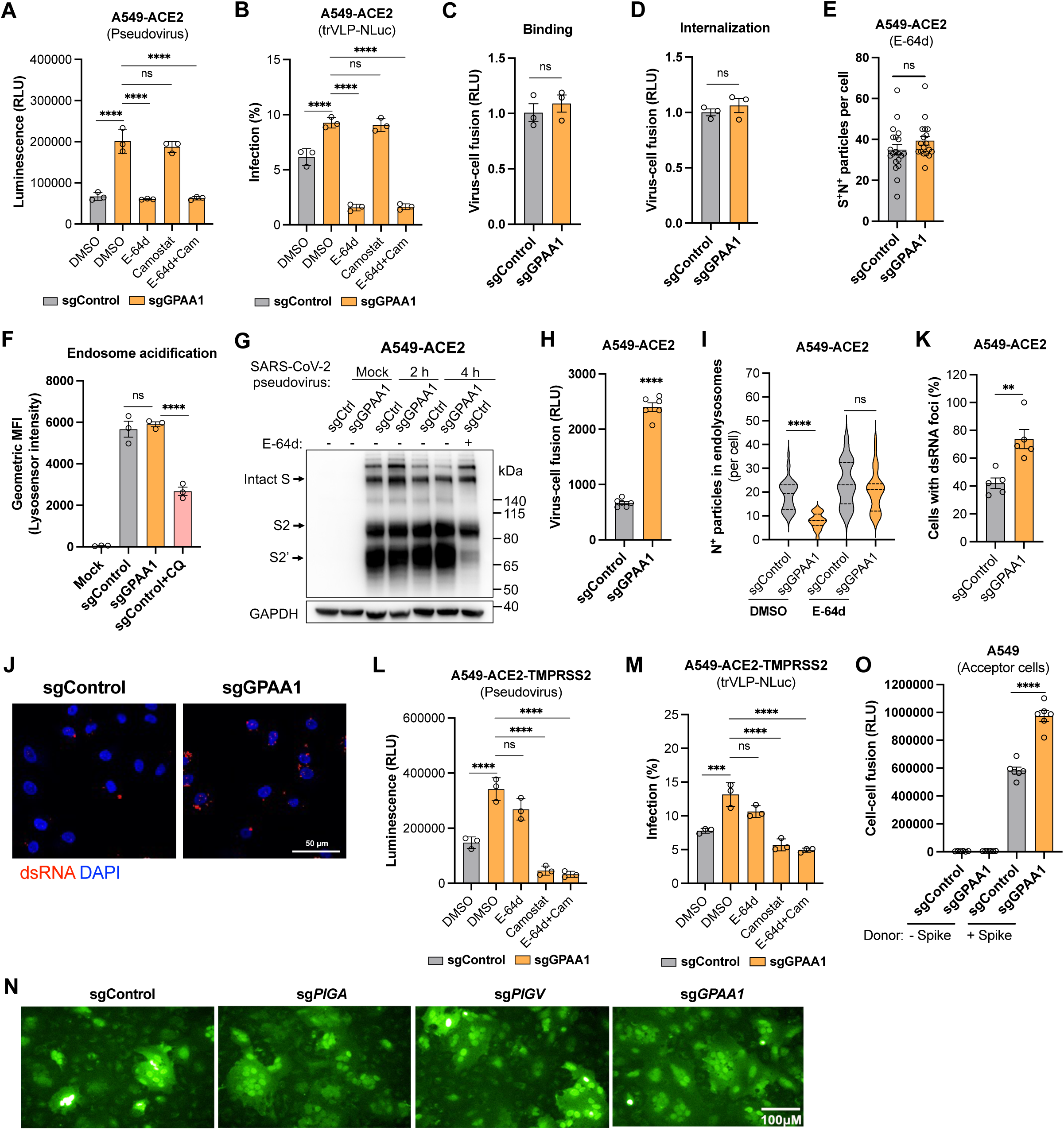
GPI biosynthesis genes inhibit coronavirus entry by disrupting spike-mediated endolysosomal and plasma membrane fusion. **A-B.** Inhibition of endosomal entry. Control and *GPAA1*-knockout A549-ACE cells were infected with MLV-based SARS-CoV-2 pseudovirus (30 μL, 36 h) **(A)** or single-round trVLP-NLuc (MOI 0.5, 24 h) **(B)** particles in the presence of cysteine protease inhibitor E-64d (aloxistatin) (100 μM) and/or serine protease inhibitor camostat mesylate (100 μM), and luciferase activity was measured. **C-D.** Virus binding **(C)** and internalization **(D)** assays. Control and *GPAA1*-knockout A549-ACE cells were incubated with SARS-CoV-2 trVLP-NLuc particles (MOI 5) on ice for binding for 45 min, or then shifted to 37°C for internalization for 45 min. The relative amount of bound or internalized virions was normalized to internal control GAPDH, and viral RNA in control cells was normalized to 1. **E.** Trafficking of SARS-CoV-2 trVLP-Nluc particles to the late endosomes or lysosomes. Control or *GPAA1*-knockout A549-ACE2 cells were incubated with the SARS-CoV-2 trVLP-NLuc particles (MOI 5, 4h) in the presence of 25 μM E-64d. The number of spike- and nucleocapsid-positive puncta co-localized with lysosome marker LAMP1 per cell was counted, and 18-20 cells from 5 fields were analyzed. **F.** Quantification of fluorescence intensities of LysoSensor Green dye in control or *GPAA1*-knockout A549-ACE2 cells in the presence or absence of chloroquine (CQ; 20 μM). **G.** Cleavage of the spike protein. Control or *GPAA1*-knockout A549-ACE2 cells were incubated with MLV-based SARS-CoV-2 pseudovirus particles in the presence DMSO or E-64d (25 μM) for 2 or 4 h, followed by western blotting analysis with anti-spike S2 antibody. **H.** Split-NanoLuc luciferase-based virus-cell fusion assay. Control or *GPAA1*-knockout A549-ACE2 cells expressing the LgBiT fragment were incubated with MLV-based SARS-CoV-2 pseudovirus containing the HiBiT fragment. Virion fusion with endolysosomes was assessed by measuring the reconstituted NanoLuc luciferase activity. **I.** Quantification of SARS-CoV-2 particles in the endolysosomes. Control or *GPAA1*-knockout A549-ACE2 cells were infected with authentic SARS-CoV-2 original strain (MOI 5, 4 h) in the presence DMSO or E-64d (25 μM). Cells were stained for confocal analysis of SARS-CoV-2 particles (N-positive) co-localized with lysosome marker LAMP1. 25-38 cells from 5 fields were analyzed. **J-K.** Quantification of dsRNA in the cytoplasm. In addition to analyzing viral particles as described in **(I)**, cells were stained with J2 antibody to detect the dsRNA foci. The representative images were visualized **(J)** and the percentage of dsRNA-positive cells per field were quantified **(K)**. 63-68 cells from 5 fields were analyzed. **L-M.** Inhibition of plasma membrane entry. Control and *GPAA1*-knockout A549-ACE cells expressing the TMPRSS2 were infected with MLV-based SARS-CoV-2 pseudovirus (30 μL, 36 h) **(L)** or single-round trVLP-NLuc (MOI 0.5, 24 h) **(M)** particles in the presence of E-64d (100 μM) and/or camostat mesylate (100 μM), and luciferase activity was measured. **N.** Representative images showing the syncytia formation after co-culture of gene-knockout A549-ACE2 cells expressing GFP1 fragment and 293T cells expressing both GFP2 fragment and SARS-CoV-2 spike protein. Scale bar, 100 μm. **O.** Split-NanoLuc luciferase-based cell-cell fusion assay. Control or *GPAA1*-knockout A549 (acceptor) cells expressing the LgBiT fragment were incubated with 293T (donor) cells expressing both HiBiT fragment and spike protein. Spike-induced cell-cell fusion was assessed by measuring the reconstituted NanoLuc luciferase activity. Data shown are from at least three independent experiments. A-B, F, I, L-M, O, one-way ANOVA with Dunnett’s test; C-E, H, K, unpaired t test; mean ± s.d.; *P < 0.05; **P < 0.01; ***, P < 0.001; ****P < 0.0001; ns, not significant.

To further confirm virion fusion with the endolysosomes, we quantified the co-localization of SARS-CoV-2 trVLP-NLuc particles (N-positive) with LAMP1. We detected fewer co-localized particles in GPAA1 knockout cells compared to controls. When the E-64d inhibitor was used to block the fusion, no significant difference in co-localization was observed between control and knockout cells **(Fig. 4I**, and **Supplementary Fig. 6)**. Additionally, we detected more double-stranded RNA (dsRNA) foci, the replication intermediates, in *GPAA1*-knockout cells than in controls **(Fig. 4J** and **K)**. These results suggest that virion fusion with endolysosomes is enhanced in *GPAA1*-knockout cells, leading to increased genomic RNA release into the cytosol to initiate replication.

### GPI biosynthesis genes inhibit coronavirus entry by disrupting spike-mediated plasma membrane fusion

In addition to endosomal entry, SARS-CoV-2 can enter cells directly through virion fusion with the plasma membrane, which requires the serine proteases such as TMPRSS2. Since A549-ACE2 cells express minimal or no TMPRSS2, we ectopically expressed the TMPRSS2 in control or *GPAA1*-knockout cells. Cells were then infected with MLV-based SARS-CoV-2 pseudovirus or single-round trVLP-NLuc particles in the presence of E-64d and/or camostat mesylate. The use of camostat inhibitor markedly diminished the enhancement of SARS-CoV-2 infection in *GPAA1*-knockout cells, whereas E-64d had minimal effect **(Fig. 4L** and **M)**, suggesting that the GPI biosynthesis pathway also modulates coronavirus entry via plasma membrane fusion.

We then examined whether GPI biosynthesis affects spike protein–mediated membrane fusion. A split-GFP system was employed to detect cell-cell fusion, as described previously^37, 38^, where fluorescence is generated only when the two GFP fragments were reconstituted in the same cell as a result of cell fusion. We observed that knockout of *GPAA1* clearly increased SARS-CoV-2 spike-mediated cell-cell syncytia formation **(Fig. 4N)**. To quantitatively measure cell-cell fusion, we employed a modified split-NanoLuc system^26^, in which the two subunits (HiBiT and LgBiT) of NanoLuc luciferase were separately expressed in A549 acceptor and 293T donor cells, respectively. The spike protein expressed in donor cells induced fusion with acceptor cells, resulting in the reconstitution of HiBiT and LgBiT to form functional luciferase. As expected, a significant increase in luciferase activity was detected in *GPAA1*-knockout cells, indicative of enhanced spike-mediated cell-cell membrane fusion **(Fig. 4O)**.

Collectively, these results suggest that the GPI biosynthesis pathway inhibits coronavirus entry by disrupting spike protein–mediated endosomal and plasma membrane fusion.

### GPI biosynthesis pathway inhibits coronavirus infection by regulating LY6E expression

The antiviral effects of the GPI biosynthesis pathway presumably depend on the downstream GPI-APs they modify, of which over 150 are known^29^. To identify which GPI-APs exert an antiviral function against coronavirus infection, we constructed a focused CRISPR knockout library with 4 sgRNAs per gene in A549-ACE2 cells, targeting 193 known or predicted GPI-APs^39, 40^ (**Supplementary Tables 2** and **3**). These cells were then infected with SARS-CoV-2 trVLP-GFP, HCoV-OC43-mGreen, HCoV-229E-mGreen (in which the mGreenLantern reporter gene was inserted into the HCoV-229E genome in place of ns4a gene), or PEDV-GFP. Similar to the genome-wide screens (**Fig. 1**), virus-infected GFP-positive cells were sorted to enrich susceptible cells for genomic DNA extraction, sgRNA sequencing, and data analysis (**Fig. 5A**). Strikingly, we found that *LY6E* was the top enriched gene in all four coronavirus screens **(Fig. 5B)**. Among the top 10 enriched genes identified for each infection condition, *LYPD2* was also identified in all four screens **(Fig. 5C)**.

**Fig. 5.**
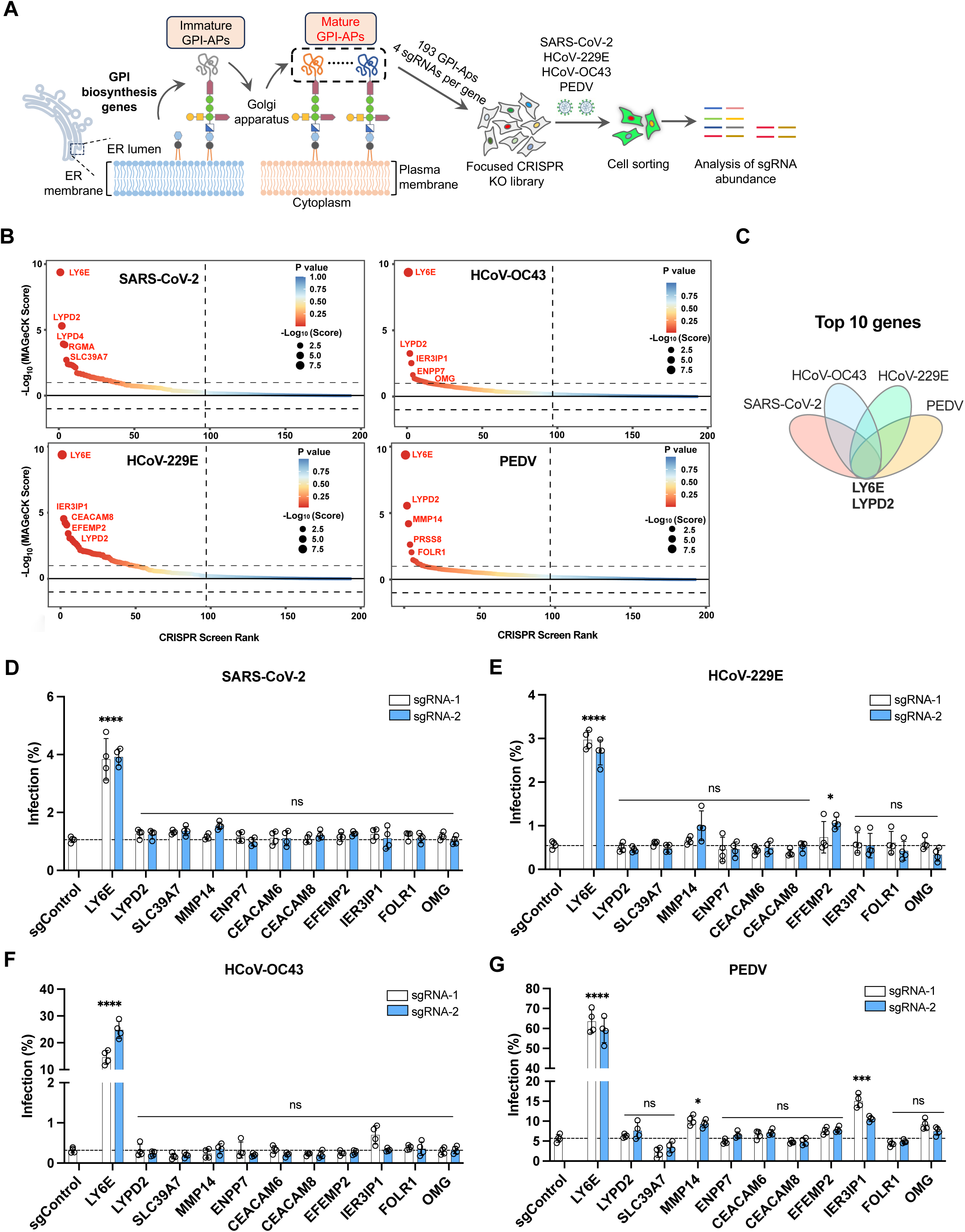
GPI biosynthesis pathway inhibits coronavirus entry by regulating LY6E expression. **A.** Schematic of focused CRISPR knockout screen of known or predicted GPI-AP genes. The sub-library of 772 sgRNAs targeting 193 known or predicted GPI-AP genes was generated, and the screens were conducted in A549-ACE2 cells infected with SARS-CoV-2 trVLP-GFP, HCoV-OC43-mGreen, HCoV-229E-mGreen, or PEDV-GFP at an MOI 0.5 for 24 h. Infected GFP-positive cells were sorted for genomic extraction, sequencing, and sgRNA analysis with MAGeCK software. **B.** The results of focused CRISPR knockout screening with four coronaviruses. The genes were ranked based on the -log_10_(MAGeCK score). **C.** Venn diagram analysis of the 10 top-ranked genes from each screen. **D-G.** Validation of the 11 genes combined from the 5 top-ranked genes from each infection screen in A549-ACE2 cells. Cells were infected with SARS-CoV-2 trVLP-Nluc (MOI 1, 15 h), HCoV-OC43 (MOI 0.5, 48 h), HCoV-229E (MOI 0.75, 48 h), or PEDV (MOI 1, 48 h), followed by flow cytometry analysis. Data shown are from four independent experiments. D-G, two-way ANOVA with Dunnett’s test; the mean of two sgRNAs was compared with the control sgRNA; mean ± s.d.; *P < 0.05; ****P < 0.0001; ns, not significant.

To validate these hits, we selected the top five enriched GPI-APs from each screen (11 genes in total) and generated A549-ACE2 knockout cell lines using two sgRNAs per gene. Upon infection with the four coronaviruses, knockout of *LY6E* led to significant increases in infection: nearly 4-fold for SARS-CoV-2, 60-fold for HCoV-OC43, 5-fold for HCoV-229E, and 10-fold for PEDV **(Fig. 5D-G)**. These findings suggest that *LY6E* is the key downstream effector mediating the antiviral activity of the GPI biosynthesis pathway.

## DISCUSSION

COVID-19 has spread globally, resulting in the most significant public health challenge since the 1918 Spanish flu^41^. Similar to other coronaviruses, SARS-CoV-2 relies on host factors to complete its life cycle, while hosts have evolved defense responses to counter viral infections^4^. Previous studies using genome-wide CRISPR screens, focused CRISPR screens, and cDNA library screens have largely focused on host proviral factors, many of which have now been identified^14, 15, 16, 17, 18, 19, 20, 24, 42, 43, 44^. However, studies of host restriction factors have been limited. Our study aimed to fill the need for a more comprehensive exploration of possible conserved host restriction factors for coronaviruses.

Encouragingly, our genome-wide CRISPR knockout screen using SARS-CoV-2, HCoV-OC43, and PEDV corroborated multiple previously identified host restriction factors. For example, *DAZAP2* and *PLSCR1* were identified in the SARS-CoV-2 screen. Additionally, many genes encoding components of GPI biosynthesis were significantly enriched in our screens. We thus focused on studying the role of these genes in regulating coronavirus infection.

GPI is a complex glycolipid composed of phosphoethanolamine, mannose, glucosamine, and phosphatidylinositol. This moiety covalently anchors proteins, known as GPI-APs, to the cell membrane, primarily in lipid raft regions^29^. Over 150 GPI-APs have been identified in mammalian cells and are essential for processes such as embryogenesis, neurogenesis, signal transduction, immune responses, and fertilization^45^. The functions of genes necessary for GPI biosynthesis thus ultimately exert their effects via the downstream GPI-APs that they modify and their role during coronavirus infection remained to be elucidated.

GPI-APs have been found to engage in the entry of many viruses. For instance, CD55/DAF acts as a receptor, facilitating the entry of hantavirus, enterovirus 70, and echovirus^46, 47, 48^. FR-α is a cofactor for the entry of Marburg and Ebola viruses^49^. NCAM-120 is an attachment factor for rabies virus^50^. In our study, we performed focused CRISPR screens targeting 193 known and predicted GPI-APs and identified LY6E as the key downstream effector of the GPI biosynthesis pathway we found in our larger genome-wide screen in restricting the infection of multiple coronaviruses. An ISG, LY6E has been shown to enhance the infection of Dengue virus, ZIKV, yellow fever virus, IAV, and VSV^51, 52^, while having dual effects on HIV-1 infection, promoting infection in CD4-high T cells and inhibiting it in CD4-low T cells^53, 54^. Recent ISG screens for SARS-CoV-2 have identified LY6E as a key host restriction factor against coronaviruses by inhibiting membrane fusion, with its conditional knockout in mice increasing murine hepatitis virus and SARS-CoV-2 infection^33, 34^. Thus, the identification of LY6E as the key GPI-AP restricting coronavirus infection could explain why the GPI biosynthesis genes, identified in our genome-wide CRISPR screen, were shown to inhibit coronavirus entry, specifically membrane fusion.

GPI biosynthesis is a complicated process that warrants further investigation. Although around 150 GPI-APs have been characterized^19^, the remaining membrane-anchored proteins regulated by the GPI biosynthesis pathway are largely unknown. Whether other GPI-APs in addition to LY6E, also play restrictive roles during coronavirus infection—perhaps affecting not only entry, but also replication, virion assembly and/or release—remains to be elucidated. However, our novel finding that GPI biosynthesis functions as a pan-coronavirus host restriction pathway provides new insights into the host defense system against virus infections and the development of host-directed antiviral therapies.

## MATERIALS and METHODS

### Cells and viruses

Vero E6 (Cell Bank of the Chinese Academy of Sciences, Shanghai, China), HEK 293T (ATCC #CRL-3216), A549 (ATCC #CCL-185), A549-ACE2^19^, HeLa (ATCC #CCL-2), Huh7, swine testicular (ST), and HRT-18 cells were all cultured at 37°C in Dulbecco’s Modified Eagle Medium supplemented with 10% fetal bovine serum (FBS), 10 mM HEPES, 1 mM sodium pyruvate, 1× non-essential amino acids, and 100 U/mL of penicillin-streptomycin. All cell lines were tested routinely and free of mycoplasma contamination.

The SARS-CoV-2 original strain (nCoV-SH01-Sfull)^19^, SARS-CoV-2 Delta variant, SARS-CoV-2 Omicron BA.1 and BA.2 variants, swine acute diarrhea syndrome coronavirus (SADS-CoV), PEDV and PEDV-GFP reporter virus (in which the ORF3 is replace by GFP), and IBV were propagated in Vero E6 cells and titrated on Vero E6 by focus-forming assay^19^. SARS-CoV-2 transcription- and replication-competent virus-like particles in which the nucleocapsid (N) gene is replaced by the reporter GFP (trVLP-GFP)^32^ or NanoLuc luciferase (trVLP-NLuc)^28^ were packaged in Vero E6 cells expressing the N protein. Porcine deltacoronavirus (PDCoV) (ST cells), HCoV-229E and HCoV-229E-mGreen reporter virus in which the ns4a is replaced by mGreenLantern (Huh7 cells), HCoV-OC43 and HCoV-OC43-mGreen reporter virus (in which the ns2a is replaced by mGreenLantern) (HRT-18 cells) were prepared and titrated similarly in their respective cell lines indicated in parentheses. All experiments involving SARS-CoV-2 authentic virus infection were performed in the biosafety level 3 facility of Fudan University.

### Genome-wide CRISPR knockout screen

The human Brunello CRISPR knockout pooled library encompassing 76,441 different sgRNAs targeting 19,114 genes^55^ was a gift from David Root and John Doench (Addgene #73178) and was packaged in HEK293T cells after co-transfection with psPAX2 and pMD2.G at a ratio of 2:2:1 using Fugene^®^HD (Promega). At 48 h post transfection, supernatants were harvested, clarified by spinning at 3,000 rpm for 15 min, and aliquoted for storage at - 80°C. For the CRISPR screens, A549-ACE2 or HeLa cells expressing the Cas9 were transduced with packaged sgRNA lentivirus library at a multiplicity of infection (MOI) of ∼0.3 by spinoculation at 1000 × g and 32°C for 30 min in 12-well plates. After selection with puromycin for approximately 7 days, cells were inoculated with SARS-CoV-2 trVLP-GFP, HCoV-OC43-mGreen, or PEDV-GFP. After infection at an MOI of 0.1 for 24 h, cells were harvested and sorted for the virus-infected reporter-positive population. Genomic DNA from both sorted cells and uninfected cells was extracted for sgRNA amplification and next-generation sequencing using an Illumina NovaSeq 6000 platform. The sgRNA sequences targeting specific genes were trimmed using the FASTX-Toolkit (http://hannonlab.cshl.edu/fastx_toolkit/) and cutadapt 1.8.1, and further analyzed for sgRNA abundance and gene ranking by a published computational tool (MAGeCK) (**Supplementary Table 1**).

### Focused CRISPR knockout screen

A total of 193 genes encoding known or predicted GPI-APs were selected for screening^39, 40^. 772 human-specific sgRNAs, with 4 sgRNAs per gene, were extracted from the human Brunello CRISPR knockout pooled library^55^, synthesized (GENEWIZ), amplified, and cloned into lentiCRISPR v2 (Addgene #52961). A549-ACE2 cells were transduced with the packaged sgRNA lentivirus sub-library as described above and then infected with SARS-CoV-2 trVLP-GFP, HCoV-OC43-mGreen, HCoV-229E-mGreen, or PEDV-GFP at an MOI of 0.1 for 24 h. The reporter-positive cells were sorted for genomic DNA extraction and sgRNA sequencing. The gene ranking was analyzed using the MAGeCK software (**Supplementary Table 2**). The KEGG pathway analysis were performed using the OmicShare tools, a online platform for data analysis (https://www.omicshare.com/tools).

### Gene editing and validation

The 50 top-ranked genes from a genome-wide knockout screen of PEDV in HeLa cells were selected for validation. For focused knockout screens of four coronaviruses in A549-ACE2 cells, 5 top-ranked genes from each screen were selected and combined (11 genes in total) for validation. Two independent sgRNAs per gene were chosen from the Brunello CRISPR knockout library and cloned into the plasmid lentiCRISPR v2 and packaged with plasmids psPAX2 and pMD2.G. A549-ACE2 or HeLa cells were transduced with lentiviruses expressing individual sgRNA and selected with puromycin for 7 days. Similarly, the GPI biosynthesis genes, PIGA, PIGV, and GPAA1 from the early, intermediate and protein-anchoring steps, were knocked out in A549-ACE2 or HeLa cells. Then, HeLa cells were infected with HCoV-OC43 (MOI 0.5, 20 h), HCoV-229E (MOI 0.75, 32 h), PEDV (MOI 1, 20 h), IBV (MOI 0.75, 20 h), PDCoV (MOI 0.25, 17 h), and A549-ACE2 cells were infected with SARS-CoV-2 trVLP-Nluc (MOI 1, 15 h), authentic SARS-CoV-2 original strain and its variants (MOI 0.5, 24 h), HCoV-OC43 (MOI 0.5, 48 h), HCoV-229E (MOI 0.75, 48 h), PEDV (MOI 1, 48 h). Cells were fixed with 4% paraformaldehyde (PFA) diluted in PBS for 30 min at room temperature, and permeabilized with 0.2% Triton x-100 in PBS for 1 h at room temperature. Cells then were subjected to immunofluorescence staining for high-content imaging or flow cytometry analysis. As for infection by viruses with reporter gene expression, cells were directly stained with 4’,6-diamidino-2-phenylindole (DAPI) without permeabilization and subjected to flow cytometry. The sgRNA sequences are listed in **Supplementary Table 3.**

### Plasmid constructs

cDNAs encoding human PIGA (NM_002641) and GPAA1 (NM_003801) were PCR-amplified from HeLa cells, fused with HA tag at their C-terminus, and subcloned into the pLV-EF1α-IRES-hygro vector. The spike gene of SARS-CoV-2 lacking the C-terminal 21 amino acids, or the full-length spike of SARS-CoV-1 was cloned into pcDNA3.1 vector. Similarly, the spike of HCoV-229E, HCoV-OC43, and PEDV were PCR-amplified and subcloned into the pCAGGS vector. The GFP1 and GFP2 fragments for the split-GFP system were synthesized (GENEWIZ), amplified, and cloned into a pCAGGS vector. The PH-Halo-LgBiT fragment was synthesized (GENEWIZ), amplified, and cloned into pLV-EF1α-IRES-Hygro. The LgBiT fragment was cloned into pCAGGS by using the synthesized PH-Halo-LgBiT as a template. The CypA-HiBiT fragment was synthesized (GENEWIZ), amplified, and cloned into pCAGGS. Lentiviruses were packaged by co-transfection with psPAX2 (Addgene #12260) and pMD2.G (Addgene #12259) using Fugene^®^HD (Promega).

### Pseudovirus packaging and infection

VSV-based pseudoviruses were produced in HEK293T cells. Cells were transfected with an expression plasmid encoding the glycoprotein gene of different viruses (SARS-CoV-2, SARS-CoV-1, HCoV-229E, HCoV-OC43, PEDV, or VSV) by Fugene^®^HD transfection reagent (Promega) for 24 h. Cells were infected with single-cycle scVSV^1′G^-Nluc-GFP^56^ in which the glycoprotein gene is deleted at an MOI of 1 for 2 h. After 3 washes, cells were maintained in culture medium in the presence of anti–VSV-G neutralizing antibody for 24 h. To package the murine leukemia retrovirus (MLV)-based pseudovirus, plasmid expressing the spike of SARS-CoV-2 was co-transfected in HEK 293T cells with the MLV-retroviral vector pMIG expressing the NanoLuc luciferase gene^19^, and plasmid expressing the MLV Gag-Pol using Fugene^®^HD transfection reagent (Promega) for 48 h. The supernatants were collected, centrifuged at 3500 rpm at 4°C for 15 min to remove cell debris, and then aliquoted for storage at −80°C. For infection, 30 µl of pseudovirus was added to each well of 96-well plates. After 12 h, the luciferase activity was determined using Nano-Glo^®^ Luciferase Assay kit (Promega #N1110) and the luminescence was recorded using a FlexStation 3 (Molecular Devices).

### Inhibition of viral entry pathways

A549-ACE2 cells with or without the TMPRSS2 expression in 96-well plate were pretreated for 1 h with cysteine protease inhibitor E-64d (aloxistatin) (100 μM) (inhibits the endosomal membrane fusion), and/or serine protease inhibitor camostat mesylate (100 μM) (inhibits the plasma membrane fusion), followed by incubation with MLV-based SARS-CoV-2 pseudovirus (30 μL, 36 h) or single-round trVLP-NLuc (MOI 0.5, 24 h) particles in the presence of inhibitors. The luciferase activity was determined using Nano-Glo^®^ Luciferase Assay kit (Promega #N1110) and the luminescence was recorded using a FlexStation 3 (Molecular Devices).

### Virus binding and internalization assay

For the virus binding assay, A549-ACE2 cells were pre-chilled on ice for 15 min followed by incubation with ice-cold SARS-CoV-2 trVLP-NLuc particles (MOI 5) on ice for 45 min. Unbound viral particles were removed by washing with ice-cold PBS three times. After the washes, cells were lysed in TRIzol reagent (Thermo fisher #15596018) for RNA extraction and qRT-PCR targeting the nsp10 gene.

For the virus internalization assay, after virus binding as described above, cells were washed with ice-cold PBS three times, followed by incubation at 37°C for 45 min. Uninternalized virions on the cell surface were removed by treating cells with 400 μg/mL protease K on ice for 45 min. After washing with ice-cold PBS three times, cells were lysed in TRIzol reagent for RNA extraction and qRT-PCR targeting the nsp10 gene. The relative amount of bound or internalized virions was normalized to the internal control GAPDH.

Viral and host mRNAs were determined using the One Step PrimeScript™ RT-PCR Kit (TaKaRa #RR064B) on a CFX Connect Real-Time System (Bio-Rad) instrument. Relative gene expression was calculated relative to GAPDH. Primers used for qRT-PCR are listed in **Supplementary Table 3.**

### Virion trafficking assay

The experiments were conducted as described previously^26^. Control or *GPAA1*-knockout A549-ACE2 cells seeded on coverslips were pretreated with 25 μM of E-64d, a cathepsin L (CTSL) proteinase inhibitor. One hour later, cells were inoculated with SARS-CoV-2 trVLP-Nluc at an MOI of 5. The inhibitor E-64d was maintained in the medium during the infection. At 4 h post infection, cells were washed twice with PBS, fixed with 4% PFA for 10 min and then permeabilized with 0.1% saponin for 10 min. Cells were blocked with 5% bovine serum albumin in PBS for 1 h and incubated with primary at 4°C overnight. After three washes, cells were incubated with the secondary antibodies for 2 h at room temperature, followed by staining with DAPI. The antibodies used are as follows: rabbit anti–SARS-CoV-2 spike protein (Sino Biological #40591-T62, 1:1000), mouse anti-SARS-CoV-2 nucleocapsid protein (made in house; 1:1000), rabbit anti-LAMP1 (Abcam #ab24170, 1:1000), goat anti-mouse or -rabbit antibody conjugated with Alexa Fluor 555 (Thermo fisher #A-21424, 2 μg/ml) or Alexa Fluor 488 (Thermo fisher #A-11034, 2 μg/ml). Images were acquired using a confocal microscope (Leica TCS SP8), and processed using Leica Application Suite X (LAS X, v3.7.0.20979). The number of spike and nucleocapsid double-positive puncta co-localized with LAMP1 per cell was quantified, with 18-20 cells from 5 fields analyzed.

### Quantification of endosomal acidification

Control or *GPAA1*-knockout A549-ACE2 cells seeded in 96-well plates were pre-treated with or without chloroquine (CQ) (20 μM). One hour later, cells were incubated in phenol red–free DMEM containing 2 μM LysoSensor Green dye (Thermo #L7535) in the presence or absence of CQ (20 μM) for 30 min at 37°C. After two washes with PBS, cells were harvested with trypsin and fixed with 2% PFA for 10 min. Cells were subjected to flow cytometry analysis (Thermo, Attune™ NxT) and the mean fluorescence intensity (MFI) of LysoSensor analyzed using FlowJo v10.0.7.

### Western blotting of cleaved spike protein

A549-ACE2 cells in 24-well plates were incubated with MLV-based SARS-CoV-2 pseudovirus particles in the presence DMSO or E-64d (25 μM) for 2 or 4 h. Cells in plates washed twice with ice-cold PBS and lysed in RIPA buffer (Cell Signaling #9806S) with a cocktail of protease inhibitors (Sigma-Aldrich #S8830). Samples were prepared in reducing buffer (50 mM Tris, pH 6.8, 10% glycerol, 2% SDS, 0.02% [wt/vol] bromophenol blue, 100 mM DTT). After heating (95°C, 10 min), samples were electrophoresed in 10% SDS polyacrylamide gels, and proteins were transferred to PVDF membranes. Membranes were blocked with 5% non-fat dry powdered milk in TBST (100mM NaCl, 10mM Tris, pH7.6, 0.1% Tween 20) for 1 h at room temperature, and probed with the rabbit anti-SARS-CoV-2 spike S2 antibody (Sino Biological #40590-T62, 1:2000) or mouse anti-GAPDH (Proteintech #60004-1-Ig, 1:2000) at 4 °C overnight. After washing with TBST, blots were incubated with horseradish peroxidase (HRP)-conjugated Goat anti-mouse (Sigma #A4416, 1:5000) or goat anti-rabbit (Thermo fisher #31460, 1:5000) secondary antibody for 1 h at room temperature, washed again with TBST, and developed using SuperSignal West Pico or Femto chemiluminescent substrate according to the manufacturer’s instructions (Thermo fisher).

### Virus-cell fusion assay

The experiments were conducted as described previously^26^. The Gag interacting protein cyclophilin A (CypA) fused with HiBiT fragment was cloned into pCAGGS. Pseudoviruses containing CypA-HiBiT were packaged in HEK 293T cells using Fugene^®^HD transfection reagent (Promega) by co-transfecting the MLV-retroviral vector pMIG^19^ in which the target gene was replaced with mGreenLantern; a plasmid expressing MLV Gag-Pol; pCAGGS expressing SARS-CoV-2 spike protein with the deletion of C-terminal 21 amino acids; and pCAGGS expressing CypA-HiBiT. At 48 h post transfection, the supernatant was harvested, clarified by spinning at 3500 rpm for 15 min, aliquoted, and stored at -80°C. Control and *GPAA1*-knockout A549-ACE2 cells (target cells) were transduced with pLV-PH-Halo-LgBiT-hygro lentivirus to stably express the LgBiT fragment. Target cells were seeded in black, clear bottom 96-well plate for 24 h, followed by spinfection with 50 μl of pseudoviruses per well at 1,000 x g, 4°C for 30 min. The virion fusion with endolysosomal membranes leads to the reconstitution of LgBiT and HiBiT as functional NanoLuc luciferase. The luciferase activity was determined at 8 h post infection using the Nano-Glo Luciferase Assay kit (Promega #N1110), and luminescence was recorded by a FlexStation 3 (Molecular Devices).

### Cell-cell fusion assay

The cell-cell fusion assay was performed according to the methods described previously with modification^37^. For visualization of the syncytia formation after co-culture of cells, we used the split-GFP system in which GFP fluorescence is generated only when its two fragments (GFP1 and GFP2) are reconstituted in the same cell (ie, due to cell fusion). Briefly, control, *PIGA-, PIGV*-, or *GPAA1*-knockout A549-ACE2 cells (acceptor cells) were transfected with pCAGGS-GFP1 (encoding the GFP1 fragment). HEK 293T cells (donor cells) were transfected with pCAGGS-GFP2 (encoding the GFP2 fragment) together with a pCAGGS vector expressing the spike protein of SARS-CoV-2 with a C-terminal deletion of 21 amino acids. At 6 h post transfection, acceptor and donor cells were trypsinized and seeded together in 12-well plates at a ratio 1:1. After 24 h of co-culture, images were captured using an AMG microscope (EVOS M7000).

To quantify cell-cell fusion based on luciferase activity as previously reported^26^, control and *GPAA1*-knockout A549 cells (acceptor cells) were transfected with pCAGGS-LgBiT encoding the LgBiT fragment of the split-NanoLuc luciferase. HEK 293T cells (donor cells) were transfected with pCAGGS-HiBiT encoding the HiBiT fragment of split-Nanoluc, together with pCAGGS vector expressing the spike protein of SARS-CoV-2 with a C-terminal deletion of 21 amino acids. At 24 h post transfection, acceptor and donor cells were trypsinized and seeded together in black, clear bottom 96-well plates at a ratio of 1:1. The spike protein-induced cell-cell fusion leads to the reconstitution of LgBiT and HiBiT as functional NanoLuc luciferase. After 24 h of co-culture, the luciferase activity was determined using the Nano-Glo Luciferase Assay kit (Promega #N1110) and luminescence was recorded using a FlexStation 3 (Molecular Devices).

### Immunofluorescence staining and analysis

For high-content imaging analysis, virus-infected cells in 96-well plates were fixed with 4% PFA for 30 min and permeabilized with 0.2% Triton X-100 for 30 min. Cells were then incubated with mouse serum (made in house; 1:1000) against nucleocapsid protein from different coronaviruses for 2 h at room temperature. After three washes, cells were incubated with secondary goat anti-mouse IgG (H + L) conjugated with Alexa Fluor 555 (Thermo fisher #A-21424) for 1 h at room temperature, followed by staining with DAPI for an additional 20 min. For infection with reporter viruses, cells were fixed and stained directly with DAPI without permeabilization and antibody incubation. Images were collected using an Operetta High Content Imaging System (PerkinElmer) and processed using the PerkinElmer Harmony high-content analysis software v4.9 and ImageJ v2.0.0 (http://rsb.info.nih.gov/ij/).

For flow cytometry analysis, virus-infected cells were harvested with trypsin and fixed with 2% PFA for 15 min. Cells were permeabilized with 0.1% saponin in PBS for 10 min, and stained with mouse serum (made in house; 1:1000) against nucleocapsid protein from different coronaviruses for 30 min at room temperature. After washing, cells were incubated with the secondary goat anti-mouse IgG (H + L) antibody conjugated with Alexa Fluor 647 (Thermo fisher #A21235, 2 μg/mL) for 30 min at room temperature. For infection with reporter viruses, cells were fixed without permeabilization and antibody incubation. After two additional washes, cells were subjected to flow cytometry analysis (Thermo, Attune™ NxT) and data processing (FlowJo v10.0.7).

For confocal imaging of virus particles in the endolysosomes or dsRNA in the cytoplasm, A549-ACE2 cells seeded on coverslips were inoculated with authentic SARS-CoV-2 original strain (MOI 5, 4 h) in the presence of DMSO or E-64d (25 μM). The inhibitor E-64d was maintained in the medium during the infection. At 4 h post infection, cells were washed twice with PBS, fixed with 4% PFA in PBS for 30 min, permeabilized with 0.1% saponin in PBS for 10 min. Cells were then incubated with primary antibody overnight at 4°C. After three washes, cells were incubated with the secondary antibody for 2 h at room temperature, followed by staining with DAPI. The primary antibodies used are as follows: mouse anti-SARS-CoV-2 nucleocapsid protein serum (made in house; 1:1000), rabbit anti-LAMP1 (Abcam #ab24170, 1:1000), mouse anti-dsRNA antibody (J2) (Scicons #10010200). The secondary antibodies used were as follows: goat anti-mouse IgG (H + L) conjugated with Alexa Fluor 555 (Thermo fisher #A-21424), goat anti-rabbit IgG (H + L) conjugated with Alexa Fluor 488 (Thermo fisher #A-11034). Images were collected using a Leica Confocal Microscope (TCS SP8) and processed using the Leica Application Suite X (LAS X, v3.7.0.20979) and ImageJ v2.0.0 (http://rsb.info.nih.gov/ij/). The number of N-positive puncta co-localized with LAMP1 per cell was quantified, with 25-38 cells from 5 fields analyzed. The total number of 63-68 cells from 5 fields were used to analyze the percentage of dsRNA-positive cells.

### Statistical analysis

Statistical significance was assigned when P values were <0.05 using Prism Version 9 (GraphPad). Data analysis was determined by an ANOVA or unpaired t-test depending on data distribution and the number of comparison groups.

## DATA AVAILABILITY

The authors declare that all relevant data supporting the findings of this study are available within the paper and its Supplementary information. The Supplemental Data provide information for the CRISPR screens, sgRNAs, and qRT-PCR analysis. Source data are provided with this paper. Any other data from this study are available upon request.

## ACKNOWLEDGEMENTS

Grants from the National Natural Science Foundation of China (82341084, 32270163, 32041005), National Key Research and Development Program of China (2024YFC2607300 and 2020YFA0707701), Program of Shanghai Academic Research Leader (22XD1420600), Shanghai Municipal Science and Technology Major Project (ZD2021CY001), Shenzhen Medical Research Fund (SMRF No. B2302029), and Non-profit Central Research Institute Fund of Chinese Academy of Medical Sciences (2023-PT310-02) supported this work. We wish to acknowledge Xiaoqing Sun, Yao Wang, and Shen Cai at Key Laboratory of Medical Molecular Virology (MOE/NHC/CAMS), Shanghai Frontiers Science Center of Pathogenic Microorganisms and Infection, School of Basic Medical Sciences of Fudan University for their help with next-generation sequencing, flow cytometry, and imaging analysis, respectively. We thank colleagues at the Biosafety Level 3 Laboratory of Fudan University for their technical assistance.

## AUTHOR CONTRIBUTIONS

Y.M., F.F., X.M., Z.W., Y.H., Y.Z., H.F., and R.Z. performed the experiments. Y.M., F.F., and R.Z. designed the experiments. Y.W., Z.G., Y.Z., and J.Z. provided technical or material support. R.Z. provided administrative, supervision support. Y.M., F.F., Z.G., and R.Z. performed data analysis. F.F. and R.Z. wrote the initial draft of the manuscript, with the other authors contributing to editing into the final form.

## SUPPLEMENTARY FIGURES

**Supplementary Fig. 1.**
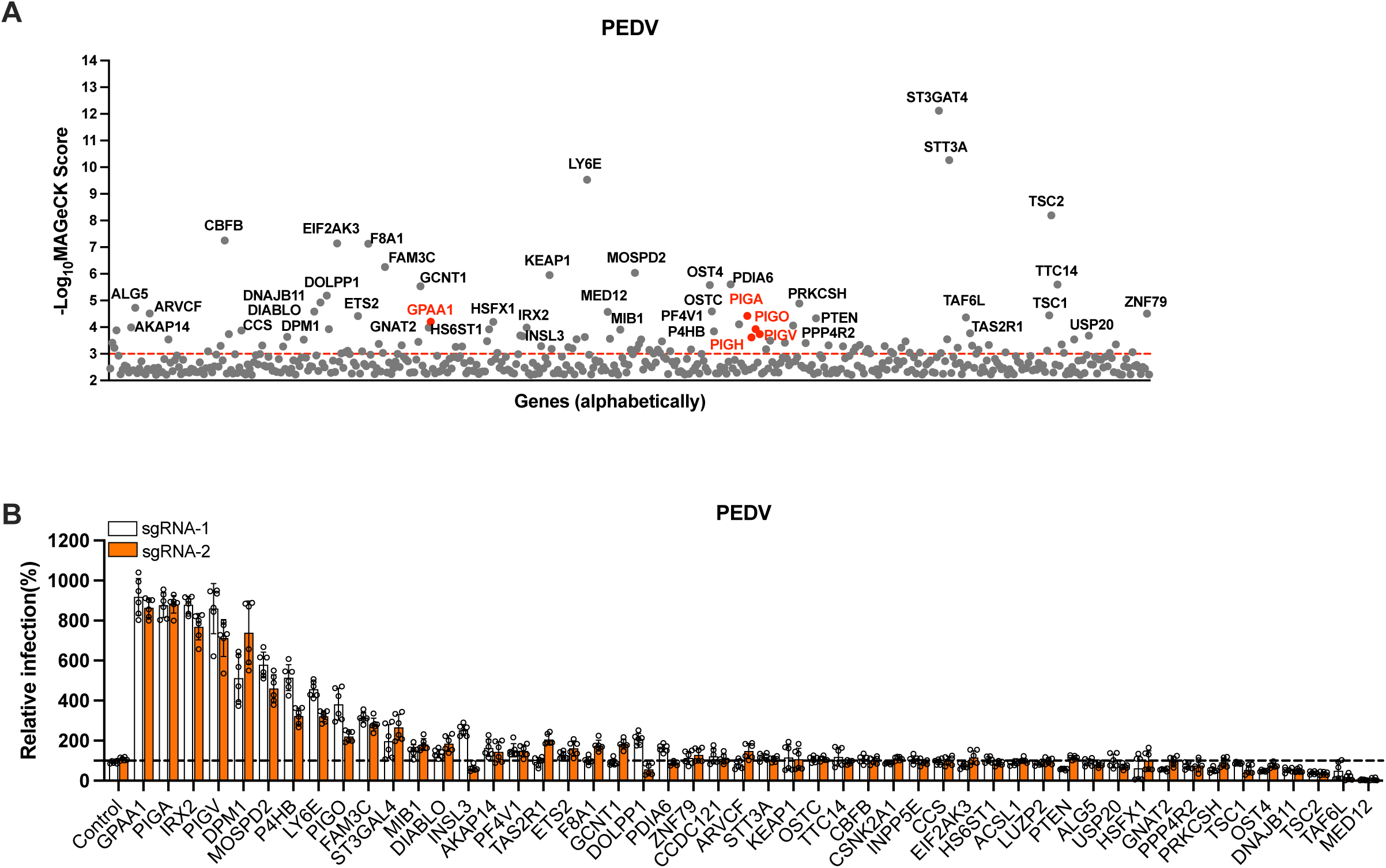
Genome-wide CRISPR/Cas9 knockout screen and validation of PEDV in HeLa cells. **A.** Genes identified from the CRISPR knockout screen in HeLa cells using PEDV-GFP. The genes were analyzed by MAGeCK software and sorted based on the -log_10_(MAGeCK score). **B.** Validation of the 50 top-ranked genes from the screen in HeLa cells. Two independent sgRNAs per gene were used, and cells were infected with PEDV (MOI 0.5, 24 h) and infection was assessed by flow cytometry. Data shown are pooled from three independent experiments, and each performed in duplicate.

**Supplementary Fig. 2.**
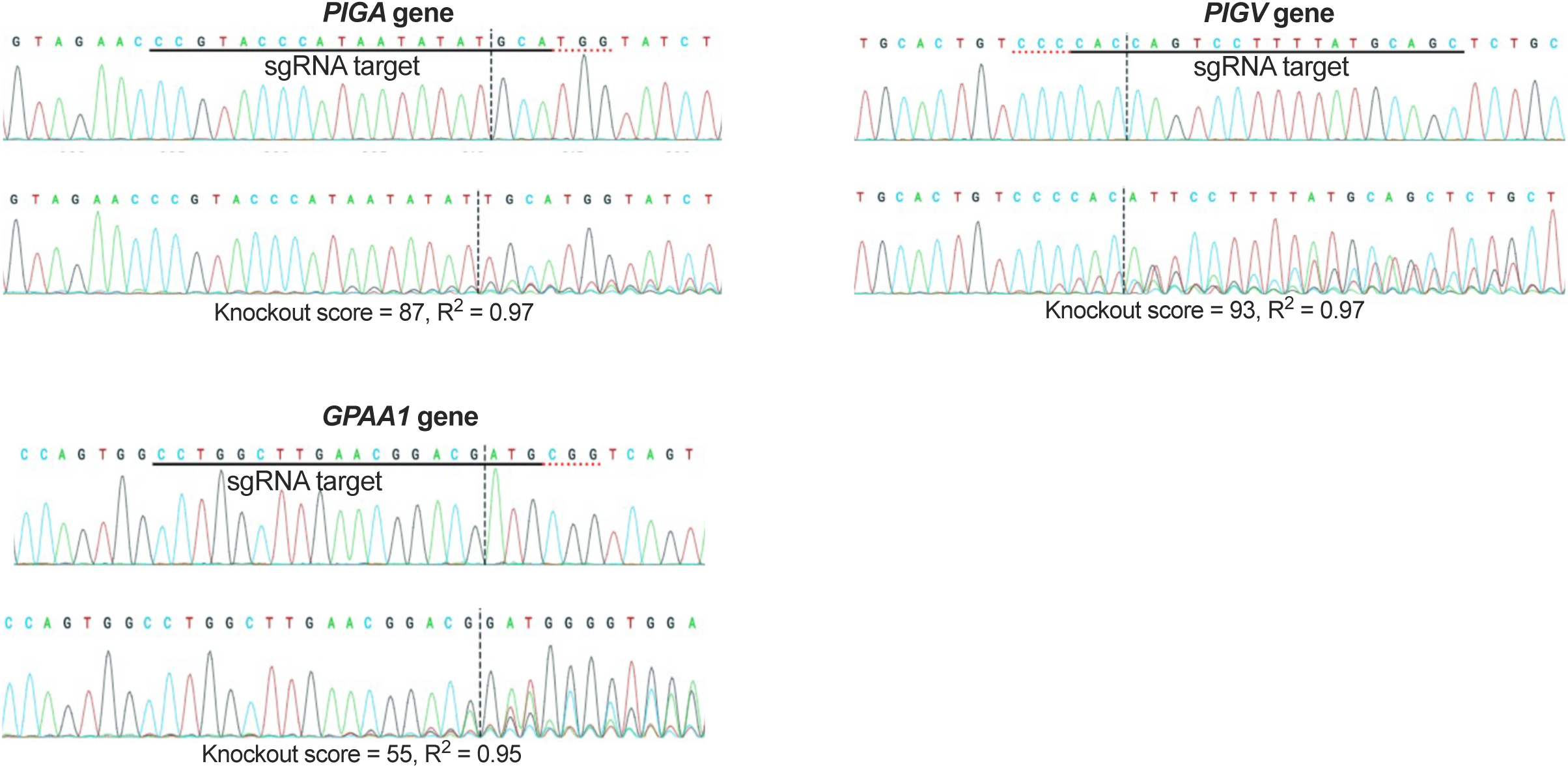
Analysis of gene knockout efficiency. **A.** The sequence traces of the gene locus of WT (upper) and *PIGA-, PIGV*-, or *GPAA1*-knockout (bottom) HeLa cells. The sgRNA target site is indicated, and knockout efficiency was determined using Inference of CRISPR Edits (ICE) analysis.

**Supplementary Fig. 3.**
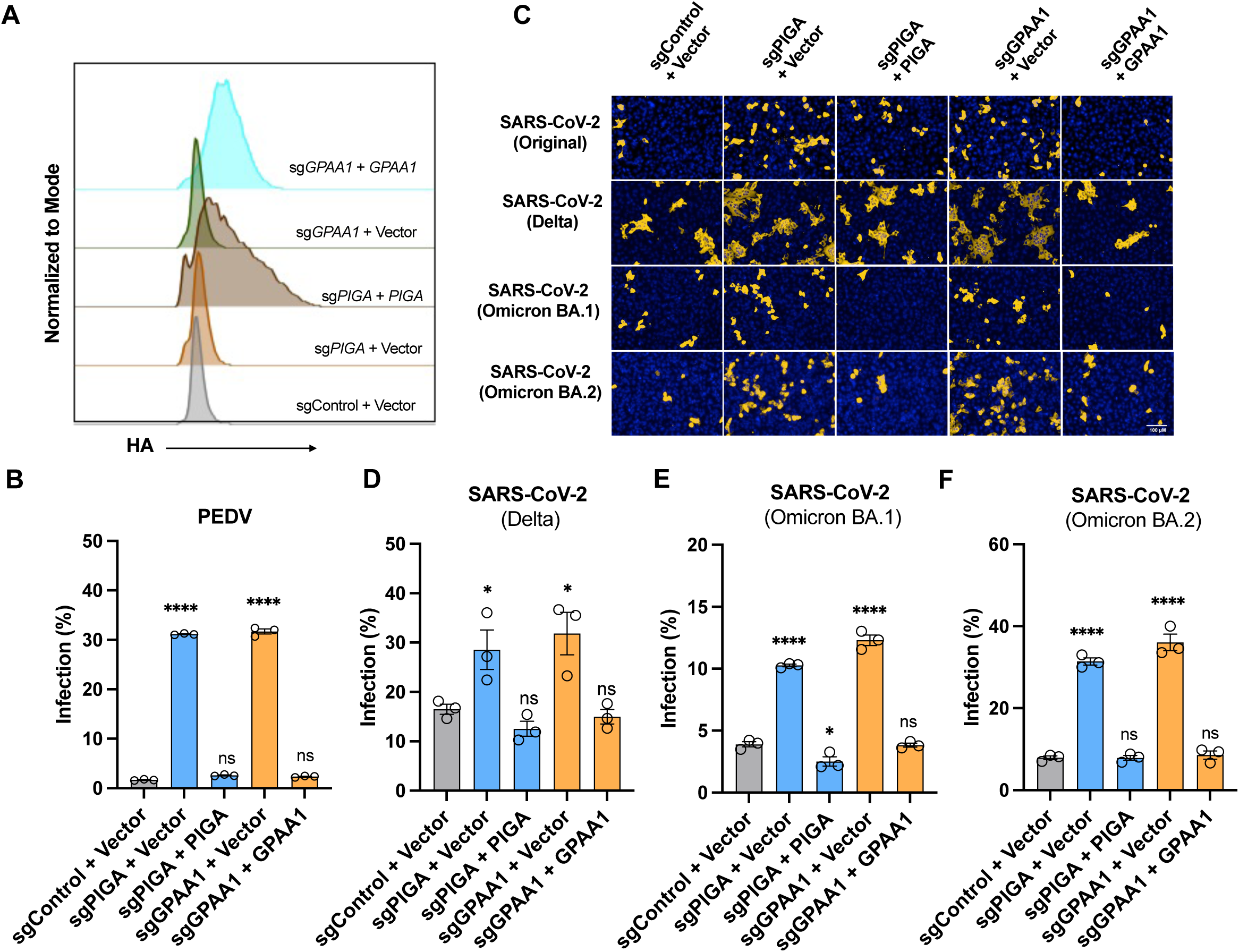
Antiviral function of GPI biosynthesis genes in knockout cells is rescued by trans-complementation. **A.** Flow cytometry analysis of *PIGA-, PIGV-,* or *GPAA1*-knockout HeLa cells trans-complemented with the respective C-terminally HA-tagged genes. Cells were trypsinized, fixed with 4% PFA, and permeabilized with anti-HA antibody. **B.** PEDV infection efficiency in trans-complemented HeLa cells (MOI 3, 20 h). **C.** Representative immunofluorescence images showing the infectivity of SARS-CoV-2 original strain and its variants in A549-ACE2 cells. Scale bar, 100 μm. **D-F.** High content imaging and quantification analysis of infection with SARS-CoV-2 Delta (MOI 0.5, 24 h) **(D)**, Omicron BA.1 (MOI 0.5, 24 h) **(E)**, or Omicron BA.2 (MOI 0.5, 24 h) **(F)** in A549-ACE2 cells. Data shown are from three independent experiments. One-way ANOVA with Dunnett’s test; mean ± s.d.; *P < 0.05; ****P < 0.0001; ns, not significant.

**Supplementary Fig. 4.**
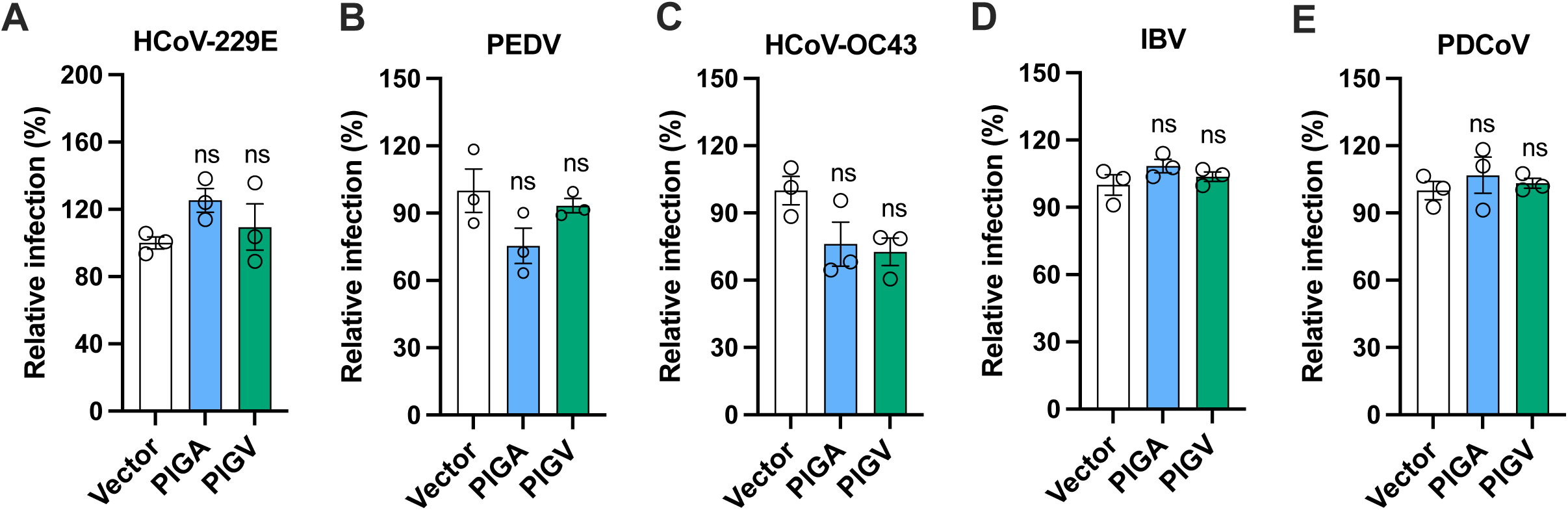
Overexpression of GPI biosynthesis genes does not enhance coronavirus infection. HeLa cells transduced with lentiviruses expressing *PIGA* or *PIGV* or a control vector were infected with HCoV-229E (MOI 0.75, 32 h), PEDV (MOI 1, 20 h), HCoV-OC43 (MOI 0.5, 20 h), IBV (MOI 0.75, 20 h), or PDCoV (MOI 0.25, 17 h). Virus infection efficiency was analyzed by flow cytometry. Data shown are from three independent experiments. Unpaired t test; mean ± s.d.; ns, not significant.

**Supplementary Fig. 5.**
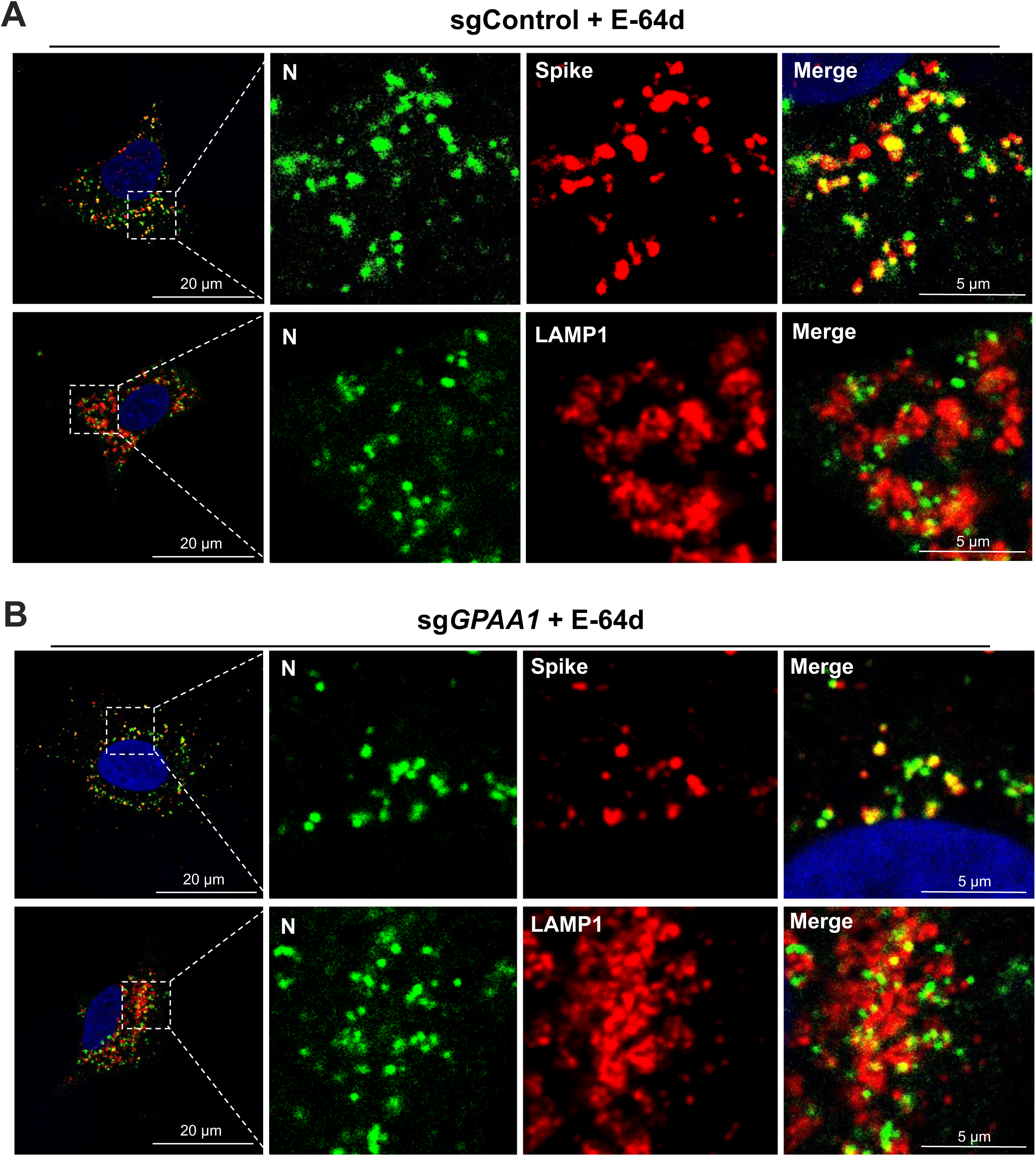
GPI biosynthesis genes do not alter virion trafficking. The control **(A)** or *GPAA1*-knockout **(B)** A549-ACE2 cells were infected with SARS-CoV-2 trVLP-Nluc (MOI 5, 4 h) in the presence of 25 μM E-64d. Cells were stained for confocal analysis of the co-localization of N and spike, or N and lysosome marker LAMP1. Representative images from three independent experiments are shown. Scale bar, 20 or 5 μm as indicated.

**Supplementary Fig. 6.**
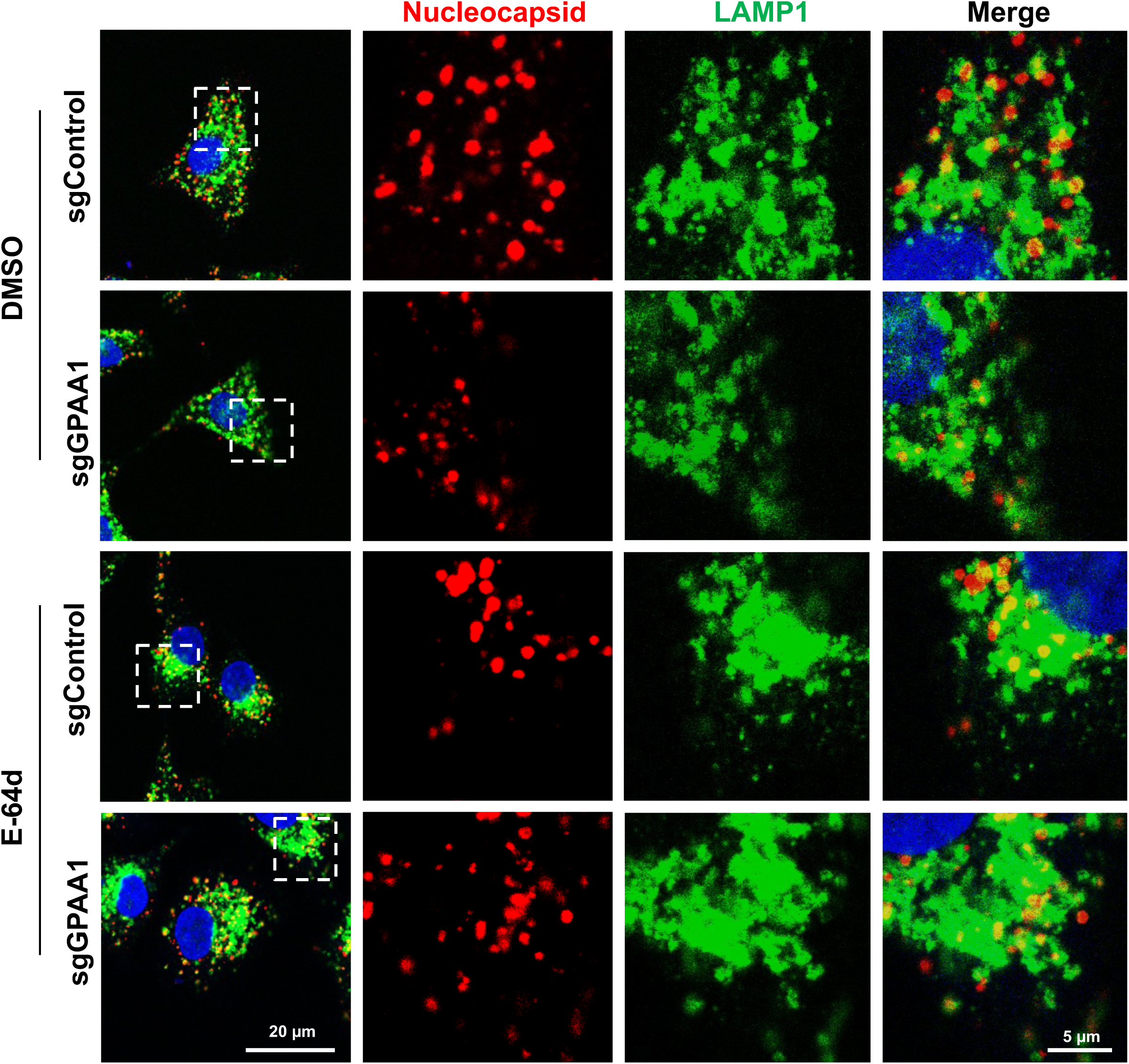
GPI biosynthesis genes disrupt the virion fusion with endolysosomes. The control or GPAA1-knockout A549-ACE2 cells were infected with authentic SARS-CoV-2 original strain (MOI 5, 4 h) in the presence of DMSO or E-64d (25 μM). Cells were stained for confocal analysis of the co-localization of N and lysosome marker LAMP1. Representative images from three independent experiments are shown. Scale bar, 20 or 5 μm as indicated.

## SUPPLEMENTARY TABLE LEGENDS

Supplementary Table 1. List of genes and scores after MAGeCK analysis of genome-wide knockout screens for three coronaviruses (see Excel file).

Supplementary Table 2. List of the known or predicted GPI-APs and scores after MAGeCK analysis of focused knockout screens for four coronaviruses (see Excel file).

Supplementary Table 3. sgRNA sequences selected for focused sub-library construction or gene validation, and primer sequences for qRT-PCR experiments (see Excel file).

